# Recurrent structural variations of lncRNA gene *CCDC26* in diffuse intrinsic pontine glioma

**DOI:** 10.1101/2021.03.14.435351

**Authors:** Lihua Zou

**Affiliations:** Northwestern University Chicago, IL, USA

## Abstract

We report recurrent somatic structural variations (SVs) involving long noncoding RNA (lncRNA) *CCDC26* in 13% of Diffuse Intrinsic Pontine Glioma (DIPG) patients. We validate our findings using whole genome sequencing data from two independent patient cohorts. *CCDC26* SVs cause increased expression of *CCDC26* gene in patients. In addition, *CCDC26* expression is associated with elevated expression of *MYC* and proliferation signature. Our findings identify *CCDC26* as a novel significantly mutated gene in DIPG and highlight the importance of structural variations in pediatric brain cancer.

## Main

Diffuse intrinsic pontine glioma (DIPG), the most frequent brainstem tumor in pediatric patients, is one of the most devastating childhood cancers, and virtually all DIPG patients die within two years after diagnosis. Current standard of care, chemotherapy followed by radiation, yields no improvements in survival. There is an unmet need for the identification of molecular mechanisms and efficacious therapeutic agents to improve treatment outcomes for DIPG patients. The discovery of somatic histone gene mutations, resulting in replacement of lysine 27 by methionine (K27M) in the encoded histone H3 proteins, in DIPG has dramatically improved our understanding of disease pathogenesis and stimulated the development of novel therapeutic approaches for the treatment of DIPG^1,2^. Past sequencing analyses of DIPG were largely focused on somatic point mutation or chromosome copy number alteration^3^. Structural variation (SV) is another class of mutations that can lead to duplication, deletion or reordering of DNA at scales ranging from single genes to entire chromosomes. The role of SVs in DIPG is poorly understood. To address this issue, we analyzed the whole genome sequences of matched tumor and normal pairs from 60 DIPG patients. These patients were from two independent cohorts: one from the CBTTC OpenDIPG project (*n* = 45) and a second from PNOS project (*n* = 15) (**Methods**). These data enable a gene-centric approach to detect SVs in DIPG.

Strikingly, we discovered recurrent SV mutations at lncRNA *CCDC26* in 13% of DIPG patients (8 out of 60) from the combined cohorts (**Fig.1a**). The role of *CCDC26* in cancer is not well understood. It was implicated in childhood acute myeloid leukemia (AML) because of altered chromosome copy numbers in AML patients^4^. We examined 8 patients with recurrent *CCDC26* SVs sample by sample. We found 6 out of 8 patients have *CCDC26* amplified through tandem duplication. To delineate critical functional region of *CCDC26*, we overlaid sequences altered by SVs at *CCDC26* locus and pinpointed a common 140kb amplicon at chr8:129,523,594-129,662,989 on hg38 reference genome (**Fig.1b**). Notably, the common amplicon is next to a germline SNP (rs4295627) associated with 1.3-fold increased risk in glioma development^5^. We observed three samples have tandem duplication breakpoints intersecting with the exon region of two neighboring genes *GSDMC* and *FAM49B* (**Fig.1b**). We searched for gene fusions involving *CCDC26* and *GSDMC* or *FAM49B* using RNA-Seq from matched samples. In patient BS_1Q524P3B, we observed a gene fusion joining transcripts from *CCDC26* exon 1 and *GSDMC* exon 7-14 (**Fig.1c**). *GSDMC* encodes Gasdermin-C, a protein coding gene which may be acting by homooligomerizing within the membrane and forming pores. We observed two patients (BS_CBMAWSAR and BS_FKQ7F6D1) displaying highly amplified DNA segments involving many breakpoints (>=100) proximal to *CCDC26*. The high copy numbers changes linked across multiple distant chromosome segments suggest ecDNA (e.g. double minute, neochromosome) as underlying structure (**Fig.1e; Fig.2c**)^6–8^. ecDNA has been reported as a mechanism which can lead to amplification of driver oncogene under selection pressure’^8^. Consistent with this, we observed co-amplified breakpoints involving *CCDC26* on chromosome 8 and *EGFR* on chromosome 7 In patient BS_CBMAWSAR (**Fig.1f**). We also observed co-amplified breakpoints involving *CCDC26* and *MYC* on chromosome 8 in patient BS_FKQ7F6D1 (**Fig.2d**).

**Figure 1.**
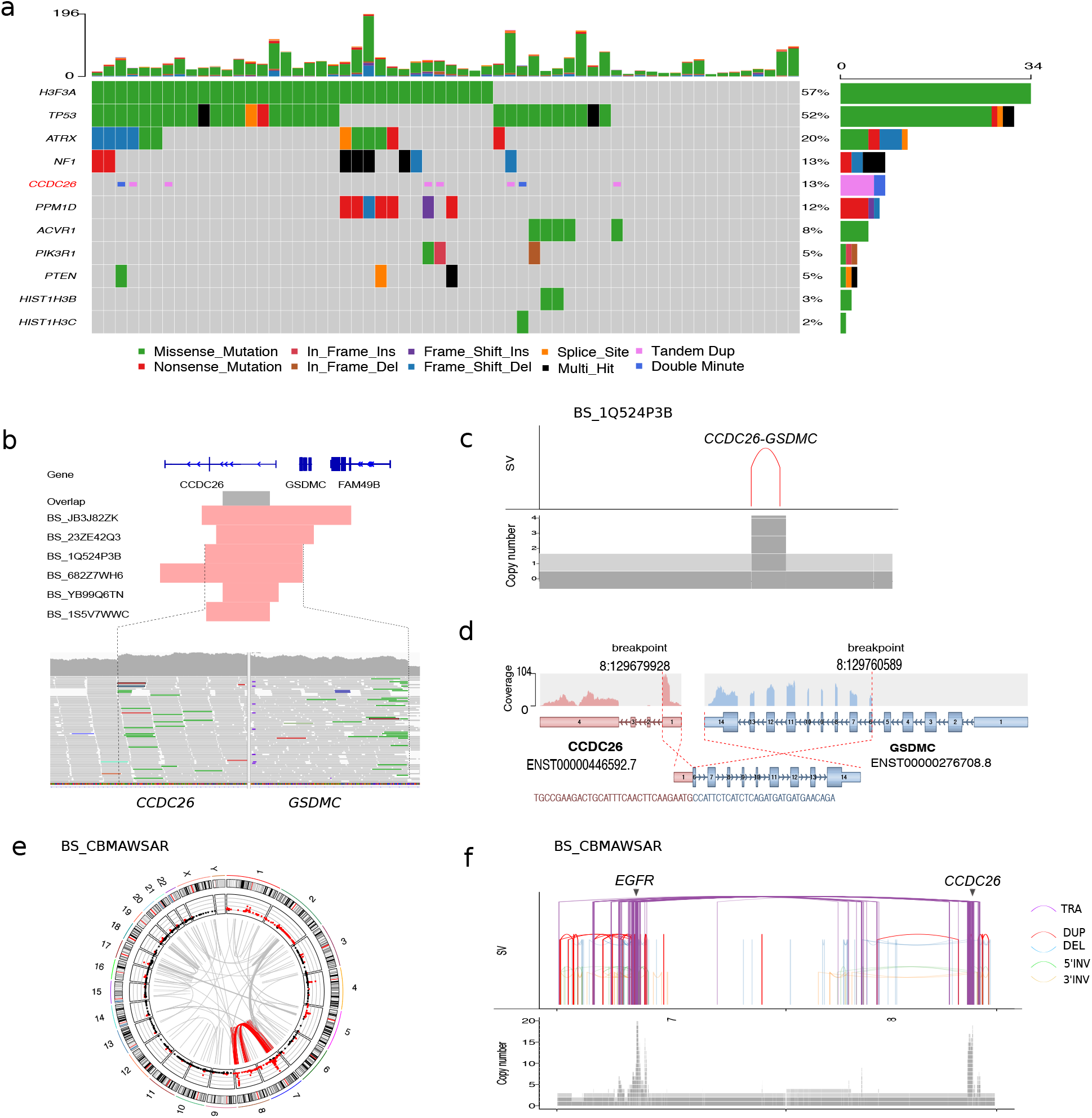
Recurrent structural variants of CCDC26 in DIPG. a) Gene-centric table showing frequency of *CCDC26* SVs along with previously reported driver mutations in DIPG in the combined cohorts; b) Common *CCDC26* amplicon (indicated by grey bar in the top panel) altered by SVs; supporting reads at breakpoints of *CCDC26* and *GSDMC* are shown in the bottom panel; c) SV and copy number changes associated with *CCDC26-GSDMC* in patient BS_1Q524P3B; d) Structure of gene fusion *CCDC26*-*GSDMC* identified in RNA-Seq of patient BS_1Q524P3B; e) Circos plots showing clustering of breakpoints at chromosome 7 and 8 in patient BS_CBMAWSAR; f) SVs (top panel) and associated high copy number changes (bottom panel) on chromosome 7 and 8 in patient BS_CBMAWSAR. Colored curves in the top panel encode different types of SVs (*red*: tandem duplication; *blue*: deletion; *green*: 5’Inversion; *orang*e: 3’Inversion; *purple*: translocation).

**Figure 2.**
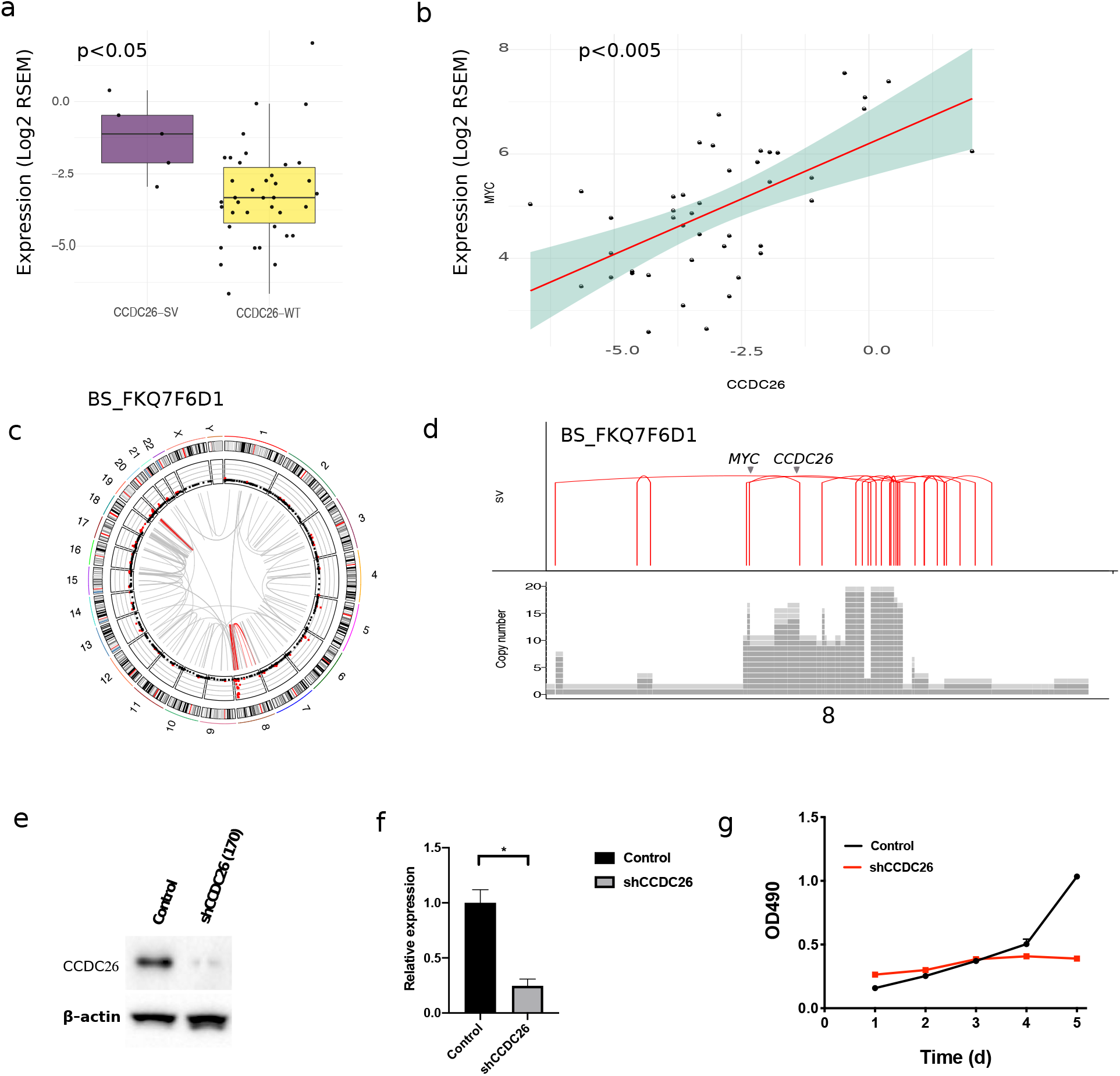
Impact of CCDC26 SVs on gene expression. a) Expression of *CCDC26* in *CCDC26*-SV vs. *CCDC26*-WT; *y-*axis indicate log2-transformed RSEM gene expression value measured by RNA-Seq; b) Correlation between *CCDC26* and *MYC* expression (Pearson correlation; p<0.005); *x-*axis and *y*-axis indicate log2-transformed RSEM gene expression value measured by RNA-Seq; c) Circos plots showing clustering of breakpoints on chromosome 8 in patient BS_FKQ7FD1; d) Detailed view of SV breakpoints and high copy number changes involving *CCDC26* and *MYC* on chromosome 8 in patient BS_FKQ7FD1.

To study the functional impact of *CCDC26* SVs, we compared *CCDC26* expression between the samples in the presence and absence of SVs (*CCDC26*-SV vs. *CCDC26*-WT) using RNA-Seq from matched samples. *CCDC26*-SV samples have significantly higher *CCDC*26 expression than *CCDC26*-WT samples (**Fig.2a**; t-test p<0.05) indicating a functional role of *CCDC26* in DIPG. To identify differential genes and pathways between *CCDC26*-SV and *CCDC26*-WT, we performed differential gene expression analysis using DESeq2^9^ and found *MYC* up-regulated in *CCDC26*-SV samples (p<0.005). We performed Gene Set Enrichment Analysis (GSEA)^10^ and found proliferation signatures and MYC targets enriched in *CCDC26*-SV samples (FDR q<0.1). In addition, *CCDC26* expression is correlated with *MYC* expression in our cohorts (**Fig.2b;** Pearson correlation p<0.005). Taken together, our findings identify *CCDC26* as a frequently mutated lncRNA gene in DIPG. Expression analysis suggests *CCDC26* is associated with *MYC* and proliferation pathways.The detailed molecular mechanism of *CCDC26* remains to be elucidated. Our findings highlight the importance of studying genome structural rearrangements in this deadly disease.

## Methods

### Cohort description

The 60 DIPG specimens used in our study are composed of radiologically diagnosed DIPG from Children’s Brain Tumor Tissue Consortium (CBTTC) and the Pediatric Pacific Neuro-oncology Consortium (PNOC). The raw whole genome sequencing and RNA-seq data can be downloaded from the Gabriella Miller Kids First Data Resource Center (KF-DRC). The CBTTC is a collaborative, multi-institutional research program dedicated to the study of childhood brain tumors. The Pacific Pediatric Neuro-Oncology Consortium (PNOC) is an international consortium dedicated to bringing new therapies to children and young adults with brain tumors. PNOC collected blood and tumor biospecimens from newly diagnosed DIPG patients as part of the clinical trial PNOC003/NCT02274987.

### Whole-genome sequencing analysis

Paired-end DNA-Seq reads were aligned to hg38 (patch release 12) reference genome using BWA-MEM^11^. Duplicates were marked using Samblaster^12^. BAMs were merged and processed using Broad’s Genome Analysis Toolkit (GATK)^13^. For WGS variant calling, Strelka2^14^ was used to call Indels and Mutect2^15^was used to call SNVs using default parameters. The final Strelka2 and Mutect2 VCFs were filtered for PASS variants for downstream analysis. For structural variant (SV) calls, Manta^16^ was used using hg38 as reference genome. Manta SV output was annotated using AnnotSV^17^. The docker image of Whole Genome Sequence Analysis Workflow can be found in the KidsFirst GitHub repository.

### Gene expression analysis

Paired-end RNA-Seq reads were aligned using ENSEMBL’s GENCODE 27 as the reference genome. Transcript- and gene-level expression values were calculated using RSEM^18^. Data normalization and differential gene expression were done by DESeq2^9^. Gene set enrichment analysis (GSEA)^10^ was used to find groups of enriched genes between different groups of samples.

### Gene fusion analysis

Gene fusions were called using Arriba^19^. Gene fusion calls from Arriba were annotated using FusionAnnotator (https://github.com/FusionAnnotator) followed by filtering of recurrent fusion artifacts and transcripts present in normal tissue using a blacklist file bundled with Arriba.

## Acknowledgements

We thank members of Children’s Brain Tumor Tissue Consortium (CBTTC) (www.cbttc.org) for their support of open access, biospecimen driven research.

